# Phylogenetic reconstruction of trait summary statistics and disparity dynamics

**DOI:** 10.64898/2026.08.16.745082

**Authors:** Gilles Didier

**Affiliations:** IMAG, Université de Montpellier, CNRS, Montpellier, France

**Keywords:** Phylogenetic comparative methods, continuous trait evolution, Brownian motion, ancestral state reconstruction, trait-distribution dynamics, empirical moments, phenotypic disparity evolution

## Abstract

Summary statistics are essential tools for understanding the overall behaviour of a collection of measurements. In a phylogenetic context, however, trait measurements are generally observed only for extant taxa and, when available, fossil taxa, while the questions of interest concern the evolution of the trait through time rather than only its static properties among the observed taxa. To investigate how the empirical mean and variance of a trait evolve through time, we derive, under Brownian evolution, their conditional distributions given trait values observed at extant or fossil tips. This provides reconstructed trajectories of both summary statistics together with a direct quantification of their uncertainty.

Among these two summaries, the empirical variance is of particular interest as a measure of past phenotypic disparity. To assess reconstructed disparity, we define the disparity level at any time as the probability that a value drawn from the conditional distribution of the empirical variance exceeds an independent value drawn from the corresponding unconditional Brownian distribution. As a dimensionless quantity with a common interpretation, the disparity level enables comparisons across times, clades, phylogenies, and datasets.

Applications to cetacean body length reveal contrasting trajectories among major subclades and suggest that much of the disparity reconstructed for the complete clade is associated with differences among these subclades. An analysis of body mass in living and fossil mammaliaforms identifies low disparity relative to the Brownian reference through most of the Mesozoic, followed by a sustained expansion around and after the K–Pg boundary. These patterns are broadly consistent with previous analyses of the same datasets, while providing a direct reconstruction of changes in the location and spread of trait distributions through time. The methods are implemented in the R package `PastMoments`.

## 1 Introduction

Reconstructing past phenotypic values is a common way to investigate how a quantitative trait has evolved through the history of a clade. A variety of approaches have been developed for this purpose, ranging from parsimony-based reconstruction to statistical methods relying on explicit stochastic models of trait evolution (Royer-Carenzi and Didier, 2016). Model-based approaches provide a probabilistic framework relating ancestral and observed trait values and, importantly, allow the uncertainty associated with ancestral reconstructions to be quantified (Martins and Hansen, 1997; Schluter et al., 1997; Garland et al., 1999). For continuous traits, many of these methods rely on Brownian motion, which remains a standard reference model in phylogenetic comparative biology (Felsenstein, 1985, 2004).

Ancestral-state reconstruction generally focuses on the value of a trait at a particular ancestral node, or, less commonly, at a specified position on a phylogeny. However, the trait values carried by the lineages present at any given time in the history of a clade form a collection whose properties are themselves informative about the evolution of the clade. As with any set of values, this collection may be usefully characterized through summary statistics rather than through its individual components. Figure 1 illustrates the summary statistics considered here, computed from a simulated Brownian trait history. At each time, the trait values carried by the lineages then present define an empirical mean and an empirical variance, whose realized trajectories are shown in the two lower panels.

**Figure 1:**
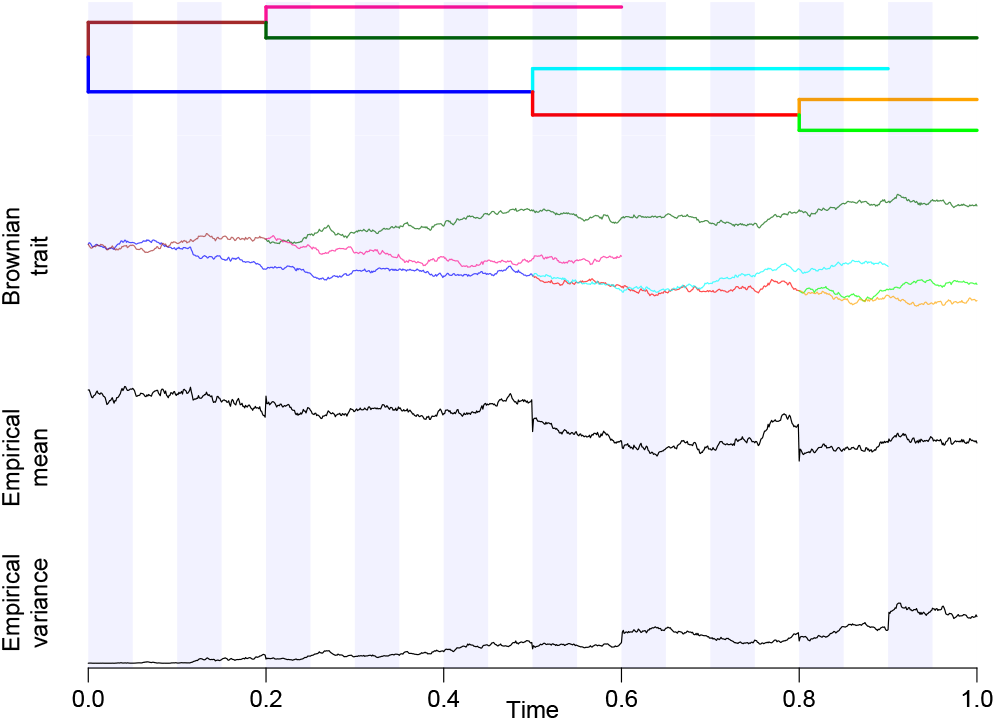
Quantities considered in this study, illustrated using a simulated Brownian trait history. Top panel: phylogenetic tree. Second panel: simulated trait evolution along its branches. Third and bottom panels: corresponding empirical mean and empirical variance, respectively, computed at each time across the lineages then present.

Of the two summary statistics considered here, the more familiar is the empirical mean, which describes the average position occupied by the clade in trait space. In the multivariate terminology commonly used in studies of morphological disparity, its natural analogue is the centroid, or mean morphology, of the occupied morphospace. To our knowledge, the temporal evolution of this centroid has been studied only in a palaeontological context, through direct comparisons of fossil assemblages from successive time intervals (Ciampaglio et al., 2001; Korn et al., 2013). By contrast, the reconstruction from tip values of the empirical mean across lineages coexisting at past times appears to have received little explicit attention, despite being relatively straightforward under the Brownian model. Under this model, conditionally on the observed tip values, the vector of trait values carried by the lineages present at any given time is Gaussian. Because the empirical mean is a linear function of this vector, its conditional distribution is itself Gaussian, with parameters determined by the phylogeny, the observed tip values, and the Brownian variance parameter.

A second fundamental summary is the empirical variance, which measures the dispersion of trait values and therefore has a direct interpretation as a measure of trait diversity or disparity. Accordingly, reconstructing the empirical variance through time provides a direct means of characterizing the temporal dynamics of phenotypic disparity. The quantification of morphological disparity and its evolution has a long history in palaeobiology and macroevolution (Wills et al., 1994; Foote, 1997; Ciampaglio et al., 2001; Guillerme et al., 2020), largely based on measurements from the fossil record.

A previous phylogenetic approach is disparity-through-time analysis (DTT, Harmon et al., 2003). At any time in the phylogeny, DTT partitions the extant taxa according to the ancestral lineages present at that time and computes the average disparity among the tip values within the resulting subclades, generally relative to the total disparity of the clade. Thus, time enters through successive phylogenetic partitions of the extant taxa, while disparity is always calculated from their present-day trait values.

Although closely related in motivation, our approach targets a different statistical object: the empirical variance among the trait values carried by lineages that actually coexisted at each past time. It reconstructs the complete conditional distribution of this collective property given the observed tip trait values, thereby quantifying its uncertainty. This reconstruction does not require individual ancestral-state estimates as an intermediate step.

More precisely, we develop a model-based framework for reconstructing the empirical mean and empirical variance of the quantitative trait values carried by the lineages present at any time in the history of a clade or, more generally, at any finite set of positions on a phylogeny. The Brownian variance parameter is estimated by restricted maximum likelihood (REML) and subsequently treated as fixed in the reconstruction. We consider the setting in which the trait value at the root is unknown and we assign it an improper flat prior. We show that integrating over this prior is equivalent to re-rooting the tree at any observed tip, fixing the trait value at the new root to its observed value, and conditioning on the values observed at the remaining tips. This equivalence reduces the reconstruction problem to standard conditioning in a multivariate Gaussian distribution. We thereby obtain the conditional distribution of the empirical mean and, by expressing the empirical variance as a Gaussian quadratic form, that of the empirical variance at any finite set of positions on the tree. We also derive explicit expressions for the conditional expectations of the empirical mean and variance, and an integral expression for that of the empirical standard deviation.

The magnitude of the empirical variance at a set of positions on a phylogeny depends on the scale of the trait, the Brownian variance parameter, and the number, temporal distribution, and phylogenetic configuration of these positions. A reference distribution is therefore needed to interpret reconstructed disparity on a common scale across analyses. For this purpose, we use the corresponding Brownian reference distribution, computed without conditioning on the tip values, for the same positions on the same phylogeny and with the same Brownian variance parameter (Didier, 2026).

We define the *disparity level* at time *t* as the probability that an independent draw from the conditional distribution of the empirical variance among the lineages present at *t* exceeds an independent draw from the corresponding unconditional Brownian reference distribution. This quantity is a probability of superiority, analogous to measures used to compare two populations (McGraw and Wong, 1992; Vargha and Delaney, 2000). It is dimensionless and has a common interpretation across phylogenies, times, clades, and Brownian variance parameters. A disparity level of 0.5 indicates no overall tendency for reconstructed disparity to be larger or smaller than under the Brownian reference, whereas values below or above 0.5 indicate lower or higher disparity relative to this reference, respectively.

Finally, we characterize the temporal form of the conditional expectations of the empirical mean and variance. Using a Brownian-bridge representation, we show that the expected empirical mean is affine between consecutive lineage event times, which correspond in practice to speciation times, fossil ages or the present, and, when all lineages of an ultrametric tree are considered, constant between consecutive lineage event times. From the structure of the conditional covariance matrix, we further show that the expected empirical variance is at most quadratic between consecutive lineage event times. Thus, between successive lineage event times, both conditional expectations follow simple polynomial forms and may exhibit instantaneous jumps at these times.

The conditional reconstruction developed here complements our recent study of the forward dynamics of these empirical moments under Brownian evolution (Didier, 2026). In that work, the trait process was considered without conditioning on observed values, and the distributions of the empirical mean and variance were studied across the lineages present through time on both fixed and random phylogenetic trees. Here, by contrast, the phylogeny and the tip values are observed, and the objective is to reconstruct the conditional distributions of the corresponding empirical summaries at past times.

We illustrate the framework with two empirical applications chosen to represent complementary phylogenetic settings. We first analyse body length in extant cetaceans (Slater et al., 2010), reconstructing the distributions of summary statistics for both the entire clade and selected major subclades. This example illustrates how the temporal dynamics reconstructed at the scale of an entire clade may combine distinct trajectories within its subclades, and allows a comparison with previous DTT results. We then analyse body mass in living and fossil mammaliaforms (Slater, 2013). The non-ultrametric phylogeny and the presence of fossils distributed throughout the history of the group illustrate how fossil observations directly constrain the reconstruction and reveal the expansion of body-size disparity around and after the K–Pg boundary.

All methods developed here are implemented in the R package `PastMoments`, which was used to generate the figures presented in this study and is available at https://github.com/gilles-didier/PastMoments.

## 2 Methods

### 2.1 Overview of the reconstruction framework

Our objective is to address the usual phylogenetic setting, in which a phylogenetic tree and trait values at its tips are observed, while both the trait value at the root and the Brownian variance parameter are unknown.

After introducing the main notation, we first consider the simpler setting in which both the root value and the Brownian variance parameter are known. In this case, Gaussian conditioning yields the distribution of the trait values at any finite set of positions on the tree conditional on the observed tip values. From this distribution, we derive the conditional distributions and expectations of their empirical mean and variance, as well as an integral expression for the conditional expectation of their empirical standard deviation. We also define their disparity level by comparing the conditional distribution of the empirical variance with its corresponding unconditional Brownian reference distribution.

We then return to the standard setting in which the root value and Brownian variance parameter are unknown. The variance parameter is estimated by REML without specifying the root value. Assigning an improper flat prior to the latter, we show that integrating it out is equivalent to rerooting the tree at an observed tip and fixing the value at the new root to its observation. The reconstruction can therefore be reduced to the known-root case treated previously.

Finally, we focus on the temporal evolution of the conditional expectations of the empirical mean and variance. After briefly recalling the Brownian-bridge representation of trait evolution along a phylogenetic branch, we show that these expectations follow simple forms between consecutive lineage event times: the expected empirical mean is affine and the expected empirical variance is at most quadratic. Both may exhibit instantaneous jumps at lineage event times.

### 2.2 Phylogenetic setting and notation

We write |*A*| for the cardinality of any finite set *A*. Let *T* be a rooted phylogenetic tree with branch lengths, and let L_*T*_ denote its set of tips. For any branch {*n, m*} of *T*, whose endpoints are the nodes *n* and *m*, we denote its length by *τ*_*n,m*_.

We refer to the node depths of *T*, i.e., the distances from the root of its nodes, as *lineage event times*. In practice, these times correspond to diversification times at internal nodes and to observation times at tips, including present-day observations and fossil occurrences.

Below, we shall consider positions on *T*, namely locations along its branches, including nodes. We represent such a position *p* by a triplet (*u, v, δ*), where (*u, v*) is an ordered pair of endpoint nodes of a branch of *T*, and where *δ* is the distance from *u* to *p* along that branch. A position of *T* may have several such representations, but this will be irrelevant in what follows. Note that this representation of positions in a tree is independent of the choice of root and therefore remains valid after re-rooting. For any node *n* of *T*, endpoint of a branch {*n, m*}, we use interchangeably *n*, (*n, m*, 0) and (*m, n, τ*_*m,n*_) for the position of *T* associated with *n*.

For any subset *D* of L_*T*_, typically a clade, and any time *t*, we define *A*^*D*^(*t*) as the set of positions of *T* that are at distance *t* from the root and lie on a path from the root to some tip in *D*.

For any two positions *p* and *q* of *T*, we denote by *t*_*p,q*_ the length of the common part of the two paths from the root to *p* and from the root to *q*. In particular, *t*_*p,p*_ is the distance from the root to *p*.

### 2.3 Known root value and Brownian variance parameter

We first determine the conditional distributions of the empirical mean and empirical variance of the trait values at a given set of positions of *T*, given the tip values, under Brownian evolution with variance parameter *σ*^2^ and root value *α*.

Let *A* = {*a*_1_, …, *a*_|*A*|_} be a non-empty finite set of positions in *T* and 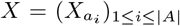 where 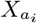 is the random variable associated with the trait value at the position *a*_*i*_ of *T*. We define the random vector 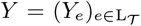 where *Y*_*e*_ is the random variable associated with the trait value at the tip *e*. The joint vector (*X, Y*) has a multivariate normal distribution with mean *α***1** and variance-covariance matrix *σ*^2^*M*, where

- for all 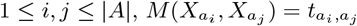,
- for all *e, f* ∈ L_*T*_, *M* (*Y*_*e*_, *Y*_*f*_) = *t*_*e,f*_,
- for all 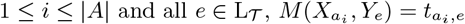.

From Anderson (2003, Theorem 2.5.1, p 35), the conditional distribution of *X* given 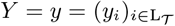 has a multivariate normal distribution with mean

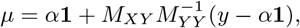

and covariance *σ*^2^*Q* with

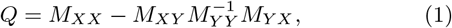

where *M*_*XX*_, *M*_*XY*_, *M*_*Y Y*_, *M*_*Y X*_ are the submatrices of *M* corresponding to the indicated rows and columns and **1** denotes the vector of ones of appropriate dimension.

#### 2.3.1 Mean

Let us now consider the empirical mean of the trait values at the positions of *A*, denoted 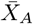, namely,

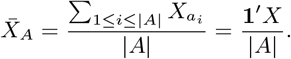

Conditionally on the tip trait values *Y* = *y*, the distribution of 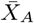 is Gaussian with mean

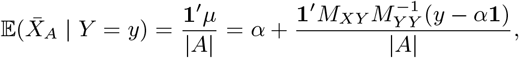

and variance

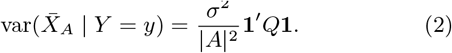

#### 2.3.2 Variance

For |*A*| ≥ 2, the empirical variance of the trait values at the positions of *A* is denoted 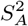 and defined as

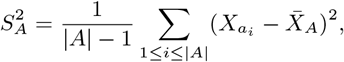

or, equivalently,

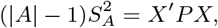

where 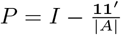, showing that 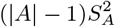 is a quadratic form in the random vector *X*.

From Mathai and Provost (1992, p. 50), the expectation of 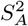 conditional on the tip trait values is

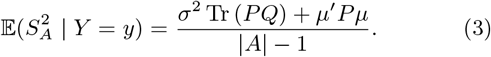

Still conditionally on the tip trait values *Y* = *y*, the distribution of 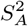 is given by

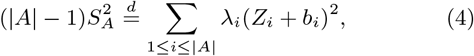

where *Z*_*i*_ are independent standard normal random variables (Mathai and Provost, 1992, p. 90). Here, assuming that *Q* is nonsingular, let

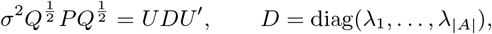

be a spectral decomposition, with *U*^*′*^*U* = *UU*^*′*^ = *I*. The vector *b* is then given by

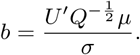

We state the nonsingular case for simplicity and refer to Mathai and Provost (1992) for the general treatment of quadratic forms in normal random vectors.

The distribution function of such Gaussian quadratic forms can be evaluated numerically using the methods of Imhof (1961) and Davies (1980); see also Duchesne and Lafaye De Micheaux (2010) for a comparison of available methods and their practical implementations.

By convention, if |*A*| = 1, the empirical variance 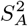 is set to zero.

#### 2.3.3 Standard deviation

Unlike the empirical variance, the standard deviation *S*_*A*_ does not admit a direct representation as a Gaussian quadratic form. However, for |*A*| ≥ 2, the conditional expectation of 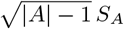 is a conditional fractional moment of 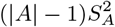, namely,

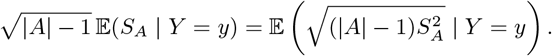

From Cressie and Borkent (1986, Prop. 5), this fractional moment can be expressed in terms of the moment generating function, or equivalently the Laplace transform, of 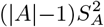 conditional on *Y* = *y*, namely

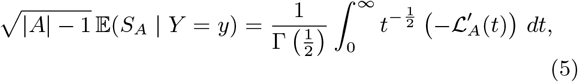

where ℒ_*A*_(*t*) is the conditional Laplace transform of 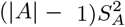 given *Y* = *y*:

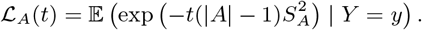

Using Equation (4) and the independence of the *Z*_*i*_, the conditional Laplace transform is

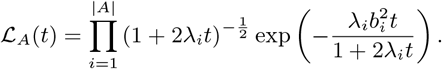

Logarithmic differentiation of ℒ_*A*_(*t*) yields

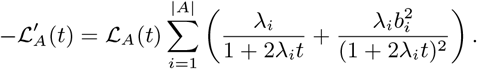

Consequently, the conditional expectation of *S*_*A*_ can be evaluated by one-dimensional numerical integration of an explicit nonnegative function (Eq. 5).

#### 2.3.4 Disparity level

Let 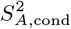 be a random variable with the conditional distribution of 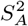 given *Y* = *y*, characterized in Equation (4), and let 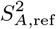 be a random variable with the unconditional distribution of the empirical variance at the same positions, under the same Brownian model on the same tree. We use the latter as a Brownian reference distribution against which reconstructed disparity is evaluated.

The unconditional distribution of the empirical variance of a Brownian trait among lineages present at a given time on a fixed phylogenetic tree was studied in Didier (2026), and the same argument applies to any finite set of positions *A*. The trait vector at these positions has mean *α***1** and covariance matrix *σ*^2^ *M*_*XX*_. Since *P* **1** = 0, the empirical variance at the positions in *A* is a central Gaussian quadratic form. Consequently,

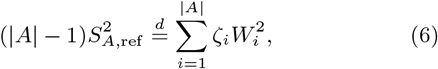

where the *W*_*i*_ are independent standard normal random variables and the *ζ*_*i*_ are the eigenvalues of 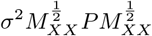.

For |*A*| ≥ 2, taking 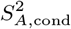 and 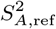 to be independent, we define the *disparity level* associated with the set of positions *A* as

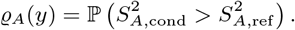

It is the probability that an independent draw from the conditional distribution of the reconstructed empirical variance exceeds an independent draw from its unconditional Brownian reference distribution. This quantity has the form of a probability of superiority used to compare two populations (McGraw and Wong, 1992; Vargha and Delaney, 2000). By convention, when |*A*| = 1, we set *ϱ*_*A*_(*y*) = 0.5.

When *A* = *A*^*D*^(*t*), we refer to 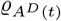 (*y*) as the disparity level at time *t* for the lineages ancestral to the tips in *D*.

In particular, if the tip values *Y* are generated under the Brownian model, then averaging the conditional distribution of 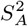 over *Y* recovers its unconditional Brownian distribution. A draw from this averaged distribution and an independent draw from the reference distribution are therefore identically distributed and, for |*A*| ≥ 2, equally likely to exceed one another because their distributions are continuous. Together with the convention adopted for |*A*| = 1, this yields

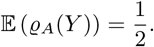

Combining Equations (4) and (6) yields

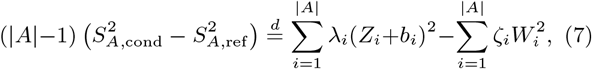

where, from the independence of 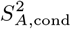 and 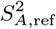, all the standard normal random variables appearing on the righthand side are independent. The difference is therefore distributed as a linear combination of independent central and noncentral chi-squared random variables, with both positive and negative coefficients. Consequently, the disparity level *ϱ*_*A*_(*y*) can be evaluated as the probability that the righthand side of Equation (7) is positive, using the same numerical methods as those used to evaluate the distribution of the empirical variance.

### 2.4 Standard phylogenetic setting

Let us now consider the standard setting of a crown tree, whose root corresponds to the first divergence, with trait values observed at the tips. Both the root value *α* and the Brownian variance parameter *σ*^2^ are unknown. The variance parameter *σ*^2^ is first estimated by REML, which does not require the root trait value *α* to be specified (Felsenstein, 2004). The resulting estimate is then substituted for *σ*^2^ in the conditional distributions of the empirical summaries computed at a set of positions on the tree.

To account for the uncertainty about the root value *α*, we assign it an improper flat prior, *π*(*x*) = 1, ∀*x* ∈ ℝ. Integrating out the root value under this prior yields a likelihood proportional, by a factor independent of the Brownian variance, to the restricted likelihood (Harville, 1974). This assumption is therefore consistent with the way in which the variance parameter is estimated. The following lemma provides a practical way to perform this integration.

#### Lemma 1.

*Let T be a rooted crown phylogenetic tree with root r, and consider a Brownian trait evolving on T with variance parameter σ*^2^. *Let A* = {*a*_1_, …, *a*_|*A*|_} *be a finite set of positions of T not containing the root r, let* 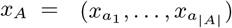, *and let f*_*T*_ (*x*_*A*_ | *x*_*r*_) *denote the joint density, evaluated at x*_*A*_, *of the trait values at the positions of A, conditional on the trait value at r being equal to x*_*r*_.

*Integrating f*_*T*_ (*x*_*A*_ | *x*_*r*_) *over x*_*r*_ *with respect to the improper flat prior with density π*(*x*) = 1, ∀*x* ∈ R, *yields, for any* 1 ≤ *i* ≤ |*A*|, *the joint density of the trait values at the positions of A \* {*a*_*i*_} *under the same Brownian model of the tree* 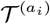, *obtained by rerooting T at a*_*i*_, *conditional on the trait value at the new root being* 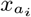. *Namely*,

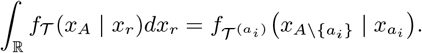

Lemma 1, together with the approach presented in Section 2.3, provides a practical way to compute, conditionally on the observed tip values, the distributions of the empirical mean and empirical variance of the trait values at any finite set of positions of a tree whose unknown root value is assigned an improper flat prior.

### 2.5 Phylogenetic Brownian bridges

In phylogenetic comparative methods, conditioning Brownian trait evolution on observed values naturally leads to Brownian bridges whose endpoint values are themselves jointly Gaussian random variables (Jhwueng, 2021; Martin and Weber, 2026).

To briefly recall the construction of such Brownian bridges, let *U* and *V* be jointly Gaussian random variables representing trait values at times 0 and *t*, respectively, and let *B* be a standard Brownian motion independent of (*U, V*). For 0 ≤ *s* ≤ *t*, define

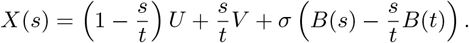

The process *X* is a Brownian bridge with random endpoints *X*(0) = *U* and *X*(*t*) = *V*. In particular,

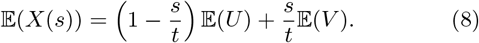

Under the setting of Section 2.3, let *T* be a stem tree, and consider a Brownian trait with known root value *α* and variance parameter *σ*^2^. Let *p* = (*u, v, δ*) be a position of *T*, where *u* is closer to the root than *v*, and where *δ* is the distance from *u* to *p* along the edge (*u, v*).

From Section 2.3, conditionally on the observed tip values *Y* = *y*, the trait values *X*_*u*_ and *X*_*v*_ are jointly Gaussian, with respective means

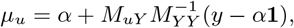

and

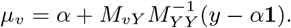

Conditionally on *X*_*u*_, *X*_*v*_, and *Y* = *y*, the trait along the edge (*u, v*) is a Brownian bridge. It follows from Equation (8) that

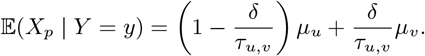

#### Lemma 2.

*Let p* = (*u, v, δ*) *be a position of T, where u is closer to the root than v, and let W* ^(*v*)^ *be the indicator vector of the set of tips descending from v, defined for all tips i by*

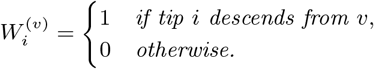

*We have*

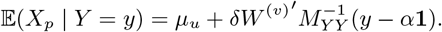

*In particular, the conditional expectation of the trait varies affinely along each edge*.

Proofs of Lemmas 1 and 2 are given in the Appendix.

### 2.6 Temporal dynamics of the conditional expectations

The preceding results characterize the conditional distribution of empirical summary statistics for an arbitrary fixed set of positions on the tree. We now consider how their conditional expectations vary through time *t* when these positions are taken to be the ancestral lineages of a given set of tips present at time *t*. Although the composition of this set of lineages changes at lineage events, the resulting temporal trajectories have a simple form between consecutive event times.

The following results are stated under the standard phylogenetic setting, that is, for a crown tree with an unknown root value.

#### Proposition 1.

*Let T be a crown phylogenetic tree, let D be a non-empty subset of its tips, and consider a Brownian trait evolving on T with variance parameter σ*^2^ *and unknown root value. Assume that the root value is assigned an improper flat prior. Conditionally on the observed tip values, the expected empirical mean of the ancestral trait values of D at time t, namely the trait values at the positions of A*^*D*^ (*t*), *is an affine function of t between consecutive lineage event times*.

*In particular, if T is ultrametric, the expected empirical mean of the trait values at all positions of T at distance t from the root is constant between successive lineage event times*.

#### Proposition 2.

*Let T be a crown phylogenetic tree, let D be a non-empty subset of its tips, and consider a Brownian trait evolving on T with variance parameter σ*^2^ *and unknown root value. Assume that the root value is assigned an improper flat prior. Conditionally on the observed tip values, the expected empirical variance of the ancestral trait values of D at time t, namely the trait values at the positions of A*^*D*^ (*t*), *is at most quadratic in t between consecutive lineage event times*.

Thus, despite the potentially complex dependence induced by conditioning on the observed tip values, the expected empirical mean and variance follow piecewise-polynomial trajectories with instantaneous jumps at lineage event times. Proofs of Propositions 1 and 2 are given in the Appendix.

## 3 Empirical analyses

The following applications illustrate both the different components of the reconstruction method and the main graphical outputs provided by the `PastMoments` package.

Following Section 2.3.4, we use the disparity level of a clade *D* at time *t*, for a tip-value vector *y*, as 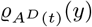, namely the disparity level associated with the ancestral lineages of *D* present at time *t*.

### 3.1 Cetacean body length

We applied our reconstruction method to the cetacean dataset of Slater et al. (2010), which combines a time-calibrated phylogeny of extant cetaceans with log-transformed average adult female body lengths. We reconstructed the summary statistics both for the complete phylogeny (Figure 2) and separately for three major clades (Figures 3 and 4).

**Figure 2:**
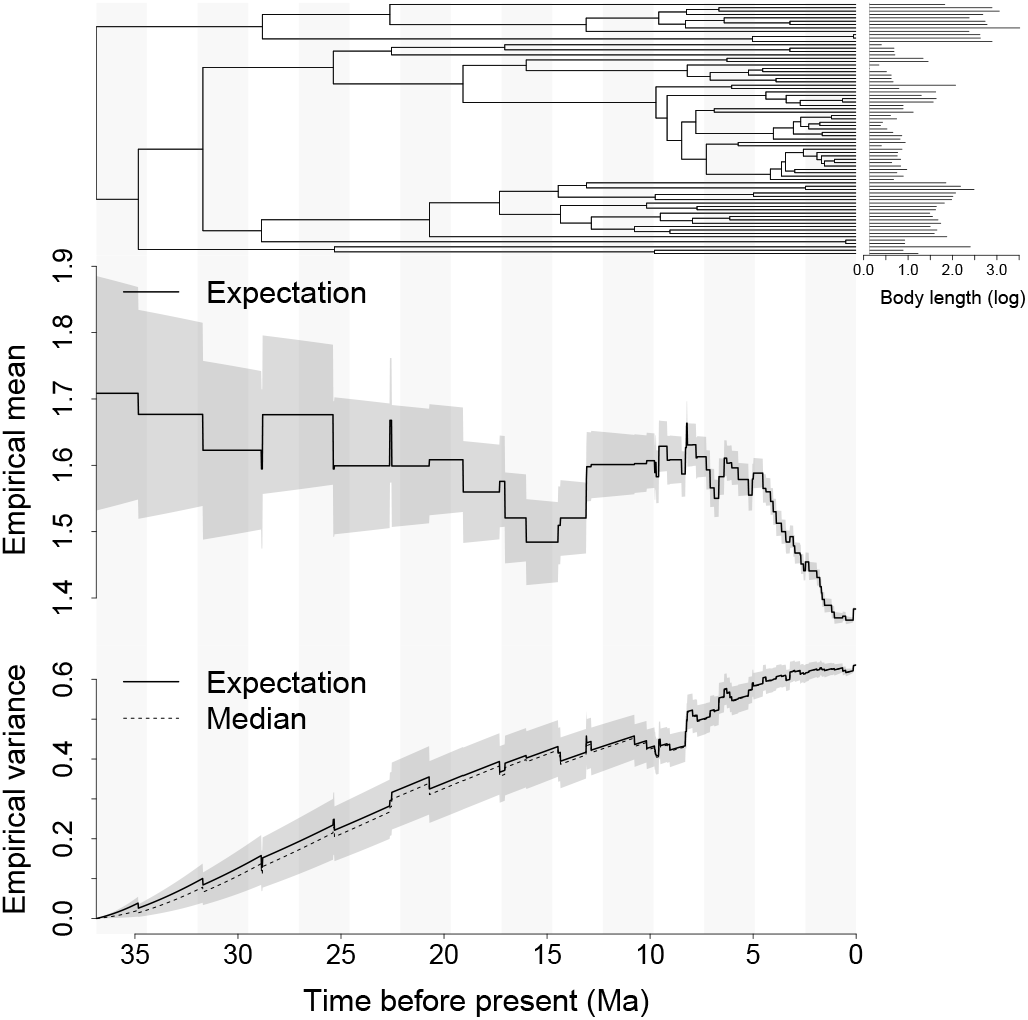
Reconstruction of summary statistics for logtransformed body length through time for all taxa in the cetacean dataset of Slater et al. (2010). The upper panel shows the timecalibrated phylogeny, with horizontal bars on the right indicating the observed log-transformed body lengths at the tips. The middle panel shows the conditional expectation of the empirical mean across all lineages present at each time. The lower panel shows the conditional expectation (solid line) and median (dashed line) of the corresponding empirical variance. Shaded bands represent the conditional interquartile ranges of the two reconstructed summary statistics.

**Figure 3:**
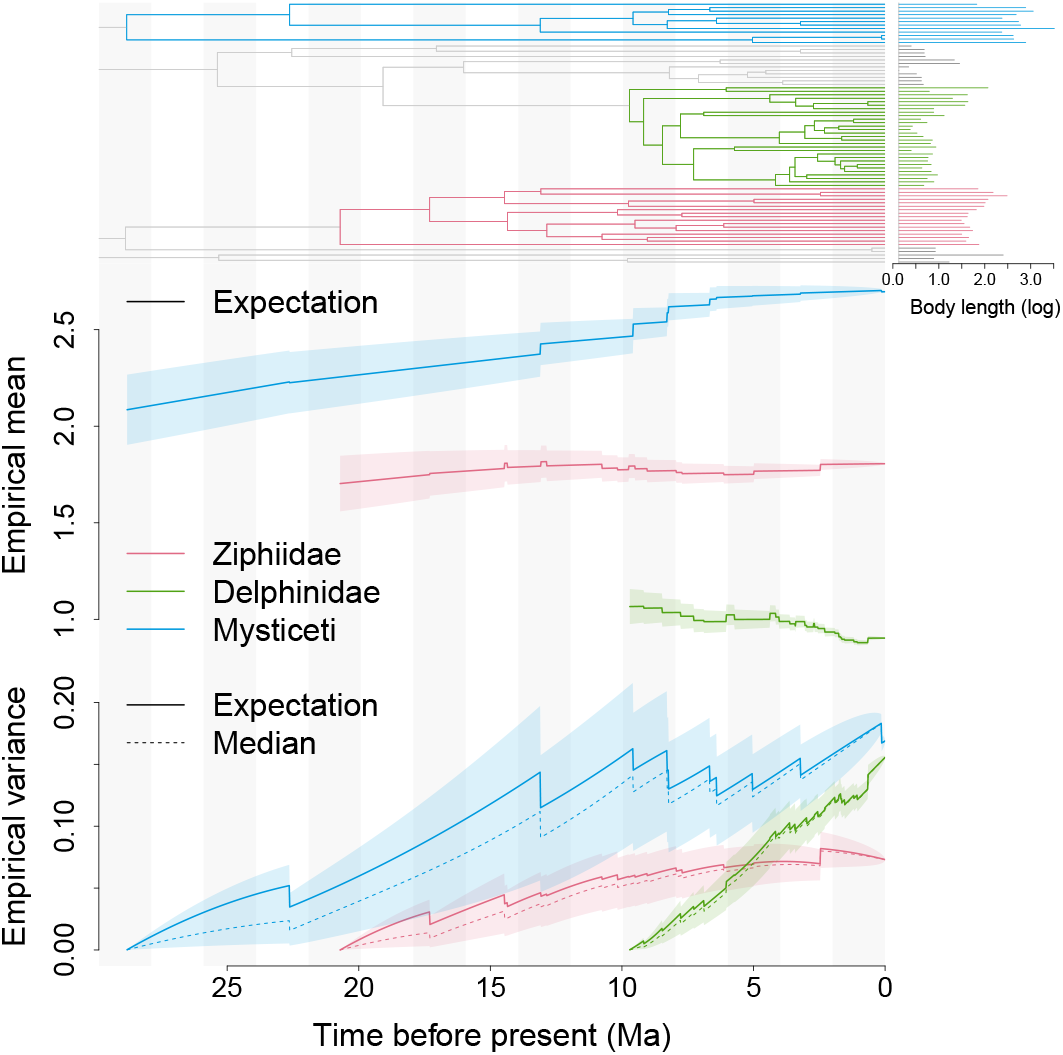
Reconstruction of summary statistics for logtransformed body length through time within three major cetacean clades in the dataset of Slater et al. (2010). The upper panel shows the time-calibrated phylogeny, with branches and observed tip values highlighted for Ziphiidae, Delphinidae, and Mysticeti. The middle panel shows the conditional expectation of the empirical mean among the lineages present within each clade at each time. The lower panel shows the conditional expectation (solid lines) and median (dashed lines) of the corresponding empirical variance. Shaded bands represent the conditional interquartile ranges of the two reconstructed summary statistics. Time is measured in millions of years before the present.

**Figure 4:**
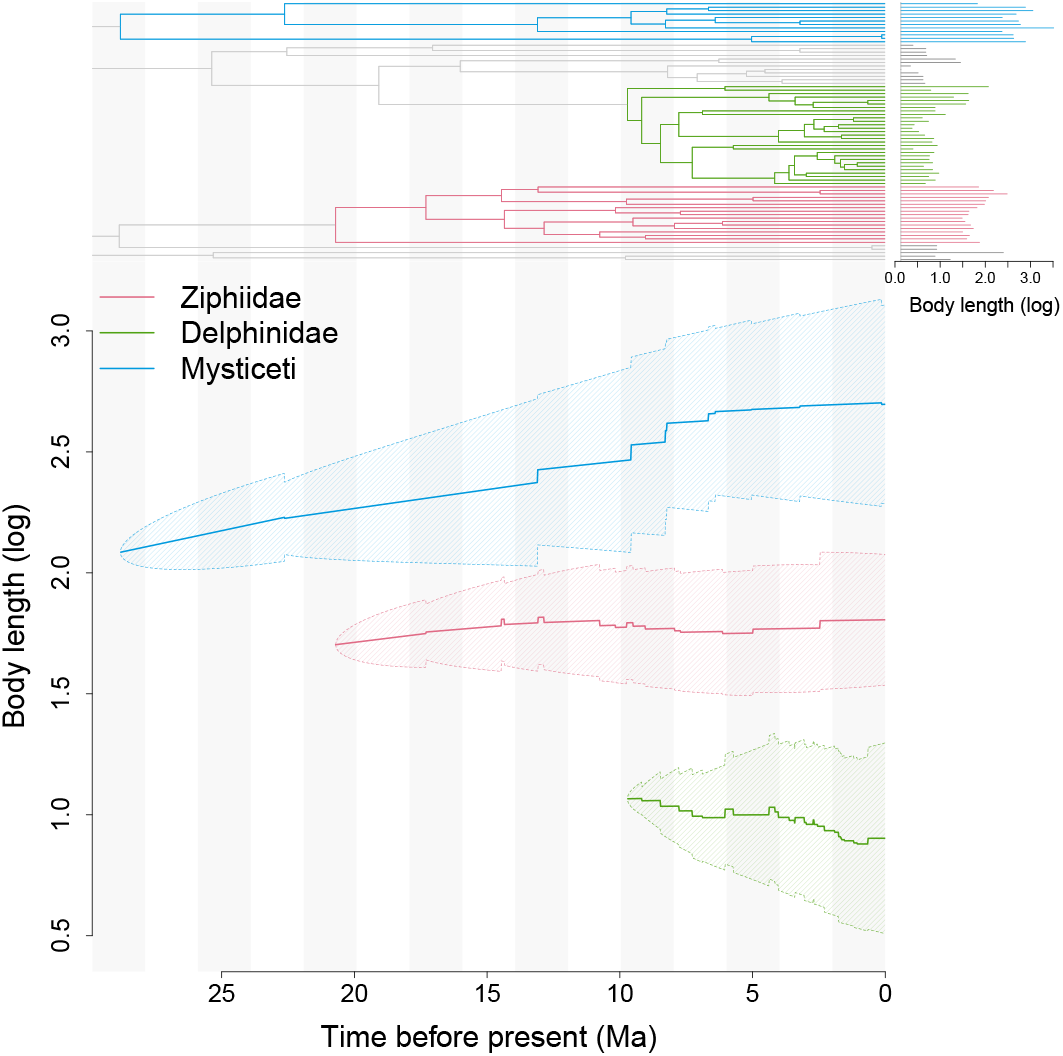
Alternative representation of reconstructed summary statistics for log-transformed body length through time within three major cetacean clades in the dataset of Slater et al. (2010). The upper panel shows the time-calibrated phylogeny, with branches and observed tip values highlighted for Ziphiidae, Delphinidae, and Mysticeti. In the lower panel, the central line for each clade shows the conditional expectation of the empirical mean among the lineages present at each time. The hatched band extends from this expectation minus the conditional expectation of the empirical standard deviation to this expectation plus the conditional expectation of the empirical standard deviation.

The first three figures provide different graphical representations of the reconstructed temporal dynamics of the empirical mean and variance and illustrate the theoretical properties established in Section 2.6. When all taxa in the dataset are considered, the conditional expectation of the empirical mean is constant between successive lineage event times, which, in the ultrametric case, correspond to speciation times and the present, as predicted by Proposition 1 (Figure 2). When the reconstruction is restricted to subclades, the conditional expectation of the empirical mean is instead affine between successive speciation times, with instantaneous jumps at these times (Figures 3 and 4).

Figures 2 and 3 also represent the conditional interquartile ranges of the reconstructed summary statistics. For the empirical mean, these intervals have zero width at the present, where all the corresponding tip values are observed, and generally widen backwards in time as ancestral trait values become less constrained by the observations.

Figure 4 provides a different representation: for each clade, the expected empirical mean is surrounded by a hatched ribbon extending from this expectation minus the expected empirical standard deviation to this expectation plus the expected empirical standard deviation, as derived in Section 2.3.3. The resulting bands do not represent uncertainty about the empirical mean, but the reconstructed spread of trait values across the lineages present at each time. They therefore provide a compact summary of the expected dynamics of trait values within the clades considered and facilitate comparison of these dynamics through time. Unlike Figures 2 and 3, however, this representation does not display the conditional uncertainty associated with the reconstructed summary statistics: the width of the hatched ribbons reflects the expected dispersion of trait values, not reconstruction uncertainty.

The variance panels in Figures 2 and 3 similarly illustrate Proposition 2. Between successive speciation times, the conditional expectation of the empirical variance is a polynomial function of degree at most two, with instantaneous jumps at these times. Moreover, at the origin of each reconstruction, only one lineage is present, so that the empirical variance is deterministically zero. At the present, all trait values are observed and the conditional distribution collapses to the observed empirical variance. Consequently, the conditional interquartile range has zero width at both endpoints, while it is generally non-zero at intermediate times.

Turning to the dataset-specific patterns, the expected empirical mean for the entire clade shows no clear directional trend through most of cetacean history, before decreasing markedly during approximately the last 5 million years (Figure 2). This relative stability at the scale of the entire clade contrasts with the distinct trajectories reconstructed within the selected subclades (Figures 3 and 4). The expected empirical mean increases substantially in Mysticeti, remains comparatively stable in Ziphiidae, and decreases in Delphinidae. Although these three subclades do not form an exhaustive partition of the cetacean clade, their contrasting trajectories show how aggregation at the scale of the entire clade can conceal marked subclade-specific trends. In particular, the recent decrease in the mean for the complete phylogeny is consistent with the radiation of relatively small-bodied Delphinidae, whose increasing number of lineages gives them progressively greater weight in the overall empirical mean.

The expected empirical variance shows a general increase towards the present, both for the entire clade and within each of the three subclades considered (Figures 2 and 3). This increase is particularly pronounced at the scale of the entire clade. Within these subclades, variance accumulates gradually over the relatively long histories of Mysticeti and Ziphiidae, although it remains substantially lower in the latter, whereas Delphinidae exhibit a rapid increase during their much more recent radiation. An increase in empirical variance is not, however, informative by itself, because variance is also expected to accumulate through time under Brownian evolution.

Comparison with the corresponding unconditional Brownian distributions reveals markedly different patterns (Figure 5). For the entire clade, the disparity level rapidly rises above 0.5 and remains close to one through most of cetacean history. The reconstructed empirical variance therefore tends to be substantially larger than expected under the unconditional Brownian model. This excess is not shared uniformly among the subclades considered. It becomes particularly strong during the recent radiation of Delphinidae, whereas the empirical variance of Ziphiidae increasingly tends to be smaller than its Brownian reference. Mysticeti remain comparatively close to the reference value, without a similarly persistent departure in either direction. Taken together with the marked separation of the subclade-specific means (Figure 4), these contrasts suggest that a substantial part of the high empirical variance reconstructed for the entire clade is associated with differences among major cetacean lineages, with an additional recent contribution from the accumulation of variance within Delphinidae.

**Figure 5:**
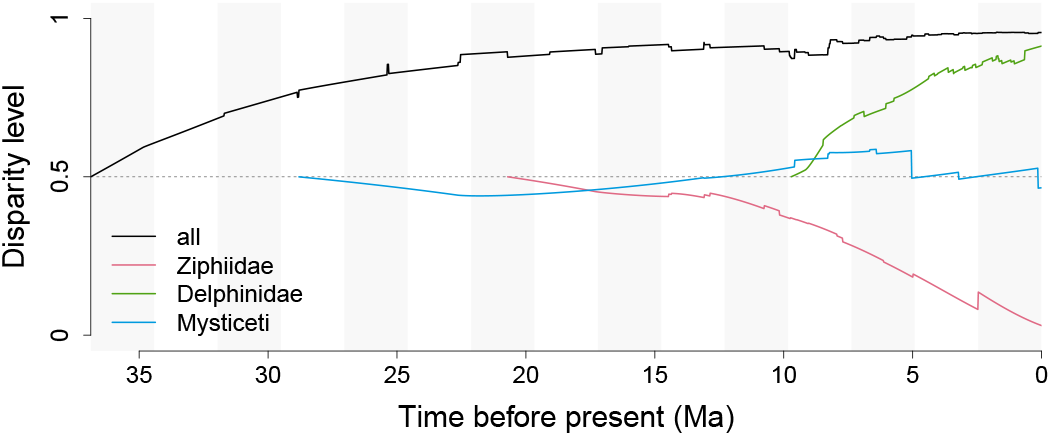
Disparity level trajectories for the entire cetacean clade and for Ziphiidae, Delphinidae, and Mysticeti. At each time, the disparity level is the probability that an independent draw from the conditional distribution of the empirical variance exceeds an independent draw from its unconditional Brownian reference distribution, both computed for the same set of contemporaneous lineages. The horizontal dashed line marks the reference value 0.5; values above and below this line indicate higher and lower disparity relative to the Brownian reference, respectively.

These reconstructed patterns are broadly consistent with the conclusion of Slater et al. (2010) that body-size niches were partitioned early among cetacean lineages and that this partitioning was broadly associated with dietary specialization. The persistent separation of the reconstructed means of Mysticeti, Ziphiidae, and Delphinidae provides a temporal view of the differentiation in body size among these major lineages. This interpretation of disparity as being primarily distributed among lineages is also consistent with the lower-than-Brownian subclade disparity reported by Slater et al. (2010), which indicates that disparity is distributed predominantly among, rather than within, subclades.

Slater et al. (2010) nevertheless identified a secondary increase in subclade disparity between approximately 11 and 6 Ma, coinciding with the radiation of several extant families and with recent dietary transitions within Delphinidae. This interval broadly coincides with the rapid accumulation of empirical variance reconstructed within Delphinidae and with the marked increase in the disparity level of this subclade.

### 3.2 Body mass in Mammaliaformes

We applied our reconstruction method to the mammaliaform dataset of Slater (2013), which combines body-mass estimates for living and fossil taxa with a time-calibrated, non-ultrametric phylogeny (Figure 6). The fossils are distributed throughout much of the Mesozoic and therefore directly constrain the reconstruction over a large part of the history of the group.

**Figure 6:**
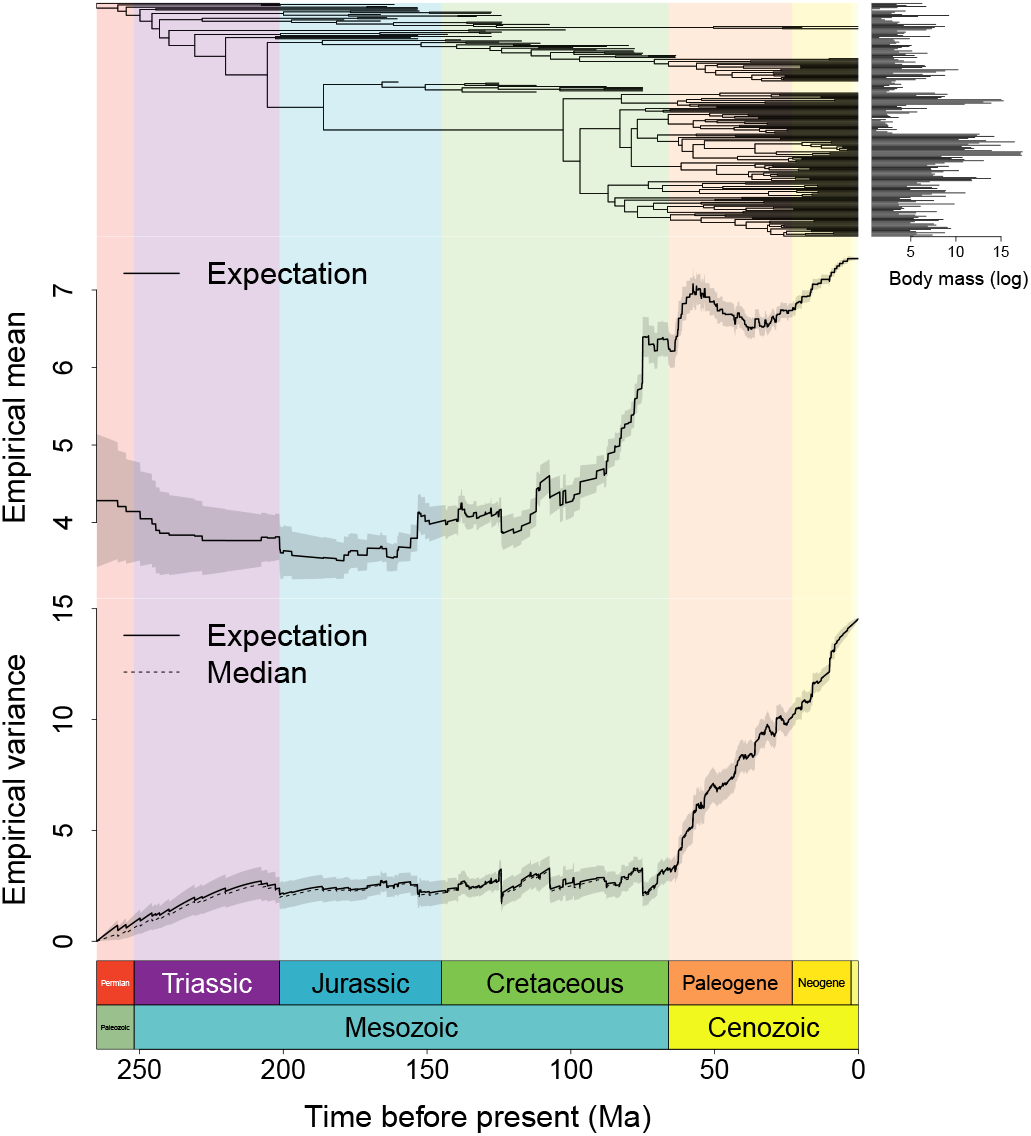
Reconstruction of summary statistics for log-transformed body mass through time for the complete set of living and fossil taxa in the mammaliaform dataset of Slater (2013). The upper panel shows the time-calibrated non-ultrametric phylogeny, with horizontal bars on the right indicating the observed log-transformed body masses at the tips. The middle panel shows the conditional expectation of the empirical mean across all lineages present at each time. The lower panel shows the conditional expectation (solid line) and median (dashed line) of the corresponding empirical variance. Shaded bands represent the conditional interquartile ranges of the two reconstructed summary statistics. Background colours indicate geological periods.

Figure 6 also illustrates the piecewise-regular behaviour established above. However, the large number of closely spaced lineage events, which here include diversification times, fossil ages, and the present, makes the individual affine or quadratic pieces less apparent than in the cetacean figures. Conditional uncertainty appears more limited than in the cetacean dataset, especially when viewed relative to the much longer temporal span of the mammaliaform phylogeny. In particular, although the interquartile range associated with the empirical mean widens backwards in time, it does so much more gradually relative to the considerably longer temporal scale of the mammaliaform phylogeny. The interquartile range associated with the empirical variance also remains comparatively narrow over most of the reconstruction. This likely reflects the stronger and temporally distributed constraints provided by the larger dataset and, in particular, by fossil observations throughout the Mesozoic.

The expected empirical mean remains low relative to its Cenozoic values and changes only moderately throughout most of the Triassic and Jurassic. It begins to increase around the middle of the Cretaceous, rises markedly thereafter, and then remains approximately stable during the last 15 Myr of the period before increasing again in the early Palaeogene. The increase thus begins well before the K–Pg boundary rather than being confined to its immediate aftermath, although the highest mean log-transformed body masses are reconstructed during the Cenozoic.

The empirical variance exhibits a different and more marked temporal pattern. Starting from zero at the root, its conditional expectation increases during the Triassic and then remains comparatively stable throughout most of the Jurassic and Cretaceous. A sustained increase begins in the latest Cretaceous, accelerates around the K–Pg boundary, and continues throughout the Cenozoic. The latter interval is thus characterized by a major expansion of body-size dispersion among contemporaneous lineages. The conditional expectation and median remain close over most of the reconstruction, while the conditional interquartile range is relatively narrow compared with the magnitude of the Cenozoic increase. The inferred pattern is therefore not primarily driven by strong asymmetry or a highly dispersed conditional distribution.

The disparity level trajectory makes the contrast between the Mesozoic and Cenozoic especially clear (Figure 7). Throughout most of the Mesozoic, the disparity level is well below 0.5 and is close to zero during much of the Jurassic and Cretaceous. It begins to rise during the Late Cretaceous, increases sharply around the K–Pg boundary, and exceeds 0.5 in the early Palaeogene. It then remains predominantly above 0.5 and increases further towards the present. Thus, relative to a homogeneous Brownian model on the same phylogeny, body-size dispersion is markedly reduced during most of the Mesozoic but becomes comparatively large during the Cenozoic.

**Figure 7:**
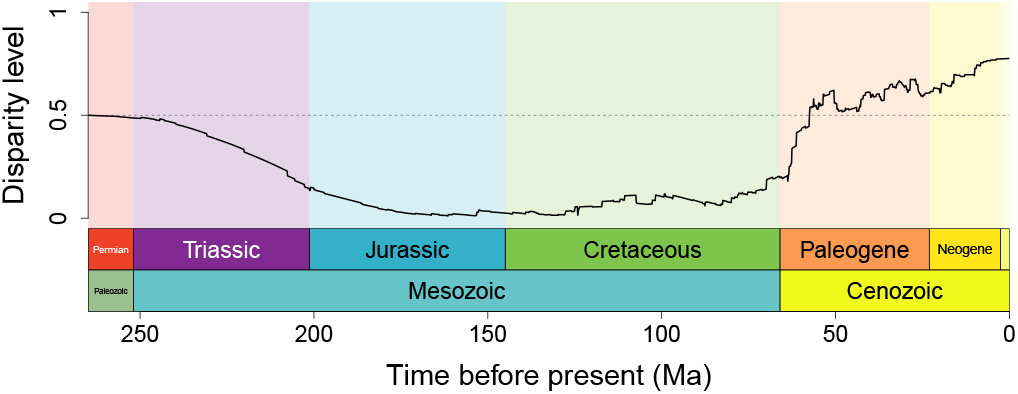
Disparity level trajectory for all living and fossil taxa in the mammaliaform dataset of Slater (2013). At each time, the disparity level is the probability that an independent draw from the conditional distribution of the empirical variance exceeds an independent draw from the corresponding unconditional Brownian reference distribution, both computed for the same set of contemporaneous lineages. The horizontal dashed line marks the reference value 0.5; values above and below this line indicate higher and lower disparity relative to the Brownian reference, respectively.

Considering the two summary statistics together, the increase in the expected empirical mean through much of the second half of the Cretaceous is not accompanied by a comparable increase in empirical variance. This combination is more consistent with a displacement of the centre of the body-size distribution towards larger values than with a simple broadening of that distribution, although the reconstructed summary statistics alone do not identify the underlying evolutionary process. By contrast, the joint increase in mean and variance during the Cenozoic indicates both an upward displacement and a broadening of the body-size distribution.

This Cenozoic pattern is qualitatively consistent with the analysis of Slater (2013), who found strongest support for a model in which body size evolved under an Ornstein– Uhlenbeck process before the K–Pg boundary and under unconstrained Brownian motion afterwards. He interpreted this transition as an ecological release from the restricted body-size distribution occupied by Mesozoic mammals, allowing body sizes to diffuse away from their physiological lower bound without requiring a directional trend. Our reconstruction does not test this evolutionary scenario directly, but recovers the same broad temporal contrast: body-size variance remains comparatively low throughout most of the Mesozoic and begins a sustained increase around the K–Pg boundary.

## 4 Discussion

The main departure of the present approach from standard ancestral-state reconstruction lies in the object being reconstructed. Rather than estimating trait values separately at individual ancestral nodes or positions, we reconstruct the conditional distributions of collective properties of the lineages present at a given time. These summaries address questions that are not naturally expressed in terms of any single ancestral state, such as the temporal displacement of the centre of a trait distribution or the accumulation of disparity among contemporaneous lineages. They are also expected to be less sensitive than individual ancestral reconstructions to local errors or stochastic fluctuations, because information is aggregated across several lineages.

The extent to which direct reconstruction of a trait summary differs from summarizing individual lineage reconstructions depends on the summary considered. Because the empirical mean is a linear function of the individual trait values, its conditional expectation is simply the average of their conditional expectations. The reconstruction of empirical moments developed here goes further by providing the full conditional Gaussian distribution of the empirical mean, thereby quantifying its uncertainty. Turning to trait dispersion, computing the empirical variance of the pointwise reconstructions accounts only for differences among the corresponding conditional means. It omits the residual conditional variances of the ancestral trait values and their conditional covariances. These terms contribute to both the expectation and the distribution of the empirical variance and are incorporated in the direct reconstruction developed here.

The present study follows a more theoretical investigation of the dynamics of empirical trait summaries under Brownian evolution without conditioning on observed tip values (Didier, 2026). Although most of that earlier work concerned random phylogenetic trees, we also obtained results for fixed trees, including characterizations of the temporal form of the empirical mean and variance that parallel Propositions 1 and 2. The temporal properties obtained here differ substantially from their unconditional counterparts. Under forward Brownian evolution, the expected empirical mean is identically equal to the initial trait value, and the non-trivial temporal result concerns its variance, which increases linearly between diversification events with slope *σ*^2^*/n* when *n* lineages are present. The expected empirical variance likewise increases linearly between events, with the universal slope *σ*^2^. After conditioning on the observed tip values, the expected empirical mean becomes data-dependent. It is affine between lineage event times and, when all lineages of an ultrametric tree are considered, constant within these intervals, although it may still jump at their boundaries. Applying the argument used in the proof of Proposition 2 to Equation (2) shows that the conditional variance of the empirical mean is at most quadratic between lineage event times. The same holds for the conditional expectation of the empirical variance, which, unlike its unconditional counterpart, need not be locally increasing.

Unlike Didier (2026), the present study is explicitly intended for application to empirical biological datasets. One of its most useful contributions in this context is the introduction of a measure of past disparity calibrated against the phylogenetic setting. Brownian motion, the standard baseline for undirected and unconstrained phenotypic evolution, provides a natural reference for this purpose, as it does in DTT analyses (Harmon et al., 2003). By comparing the conditional distribution of the reconstructed empirical variance with its unconditional Brownian distribution for the same positions on the same tree, the disparity level places reconstructed disparity on a common, dimensionless scale.

The disparity level belongs to the class of probability-of-superiority measures used to compare two populations. It summarizes the relative positions of independent draws from the complete conditional and unconditional distributions of the same empirical-variance statistic. It therefore depends on the locations, dispersions, and shapes of both distributions rather than only on their expectations or medians. It is dimensionless and has a direct interpretation: values above 0.5 indicate that the reconstructed empirical variance is more often larger than its Brownian reference than smaller, whereas values below 0.5 indicate the converse. Because it depends only on the ordering of the two draws, it is also invariant under any common strictly increasing transformation applied to them. These properties make it comparable across phylogenies, sets of positions, and Brownian variance parameters.

We also considered the relative difference between the conditional and unconditional expectations of the empirical variance, but retained the probability-of-superiority measure because it depends on the full conditional and unconditional distributions rather than only on their expectations.

The graphical representations developed here are primarily exploratory. By displaying the reconstructed distributions of past empirical summaries, they make it possible to examine jointly the displacement of the centre of the trait distribution, changes in its dispersion, uncertainty about these quantities, and their departure from the corresponding unconditional Brownian distributions. These components need not exhibit parallel dynamics. Their joint representation can therefore suggest interpretable temporal scenarios that would be difficult to identify from separate ancestral-state reconstructions or from the trajectory of a single summary statistic.

The cetacean analysis illustrates the value of applying the reconstruction at different phylogenetic scales. The relatively stable empirical mean reconstructed for the entire clade results from the aggregation of strongly contrasting trajectories within major subclades, while the high overall empirical variance reflects both variation within these subclades and the separation of their mean trait values. A global trajectory therefore need not be representative of the dynamics occurring within its constituent lineages. Conversely, comparisons among subclades can help identify whether the accumulation of disparity is broadly shared or concentrated within particular radiations.

The mammaliaform analysis shows how fossil observations, by providing information at intermediate times, can reduce reconstruction uncertainty relative to analyses based exclusively on extant taxa. The joint examination of the empirical mean and variance also provides complementary information. In this example, the increase in mean body size begins before the major expansion of variance, suggesting an initial displacement of the centre of the body-size distribution followed by a broader diversification of body sizes.

One caveat concerns the treatment of the Brownian variance parameter, which is estimated by REML and then treated as known. This approximation has no effect on the conditional expectation of the empirical mean, because the conditional mean trait values do not depend on the Brownian variance parameter. For the empirical variance, the contribution arising from differences among the conditional mean trait values is likewise unaffected, whereas the contribution of residual conditional uncertainty is proportional to the estimated variance parameter. Consequently, the reconstructed distribution of the empirical variance and the resulting disparity level do not account for uncertainty about the Brownian variance parameter. This uncertainty could be incorporated in a Bayesian extension by assigning a prior to this parameter and integrating the reconstruction over its posterior distribution, for example using MCMC. We do not pursue this extension here because the present study focuses on the analytical reconstruction of empirical moments.

The reconstruction approach presented here extends directly to other Gaussian evolutionary models when their parameters and initial-state distributions are specified. In particular, under an Ornstein–Uhlenbeck model, replacing the Brownian mean vectors and covariance matrices with their Ornstein–Uhlenbeck counterparts yields the same general results: the conditional empirical mean is Gaussian, while the empirical variance remains a noncentral Gaussian quadratic form. Its expectation and distribution, the expected empirical standard deviation, and a corresponding disparity level can therefore be computed by the same methods. Extending the full framework is less straightforward when the evolutionary parameters and initial state are unknown. In particular, the Brownian rerooting equivalence does not carry over directly, and different treatments of the root state and optimum define genuinely different models. We explored several such variants but defer their systematic investigation to future work in order to keep the present study focused on the Brownian case.

To conclude, this study builds on the idea that the evolving distribution of trait values among contemporaneous lineages can itself be treated as an object of phylogenetic reconstruction. By jointly considering changes in its location and dispersion, this perspective provides a direct view of trait-distribution dynamics through the history of a clade.

## 5 Acknowledgements

ChatGPT and Codex (OpenAI) were used for writing assistance and coding support. The author takes full responsibility for the final content.

## A Proof of Lemma 1

Let us assume that all the positions of *A* are nodes of *T*, up to adding new nodes in *T* for all positions of *A* strictly inside branches of *T*. We write **N**_*T*_ for the set of nodes of *T* and **E**_*T*_ for the set of edges of *T* oriented away from its root.

For all real values *x* and *y*, the Brownian transition density *g*_*τ*_ (*x, y*) along a branch of length *τ* is the density at *y − x* of a centered Gaussian distribution with variance *σ*^2^*τ*.

The joint density of the trait values at the nodes of *A*, conditional on the trait value at *r* being equal to *x*_*r*_ is then

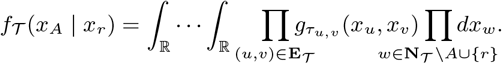

Let us now consider the tree 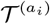 obtained by rerooting *T* at the node *a*_*i*_. We have that 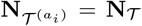 and 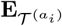 is obtained from **E**_*T*_ just by changing the orientation of the edges in the path from *r* to *a*_*i*_.

Since the Brownian transition density is symmetric, i.e., *g*_*τ*_ (*x, y*) = *g*_*τ*_ (*y, x*) for all real values *x, y* and all lengths *τ*, we have that

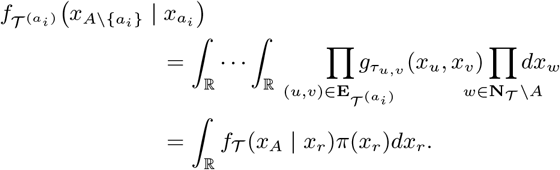

## B Proof of Lemma 2

For every tip *i* of *T*,

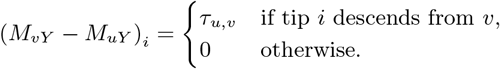

Consequently,

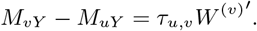

Equation (8) gives

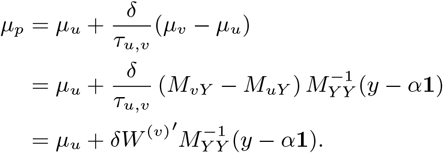

## C Proof of Proposition 1

We consider the natural direction of time and assume, without loss of generality, that the origin of time corresponds to the root *r* of the crown tree *T*. For any time *t*, let 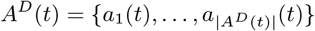. We write *a*_*i*_(*t*) = (*u*_*i*_, *v*_*i*_, *δ*_*i*_), with *u*_*i*_ closer to *r* than *v*_*i*_.

Let *t* be a time in the temporal span of *T* and let Δ ∈ ℝ be such that there is no lineage event time between *t* and *t* + Δ in *T*. It follows that |*A*^*D*^(*t* + Δ)| = |*A*^*D*^(*t*)| and, for all 1 ≤ *i* ≤ |*A*^*D*^ (*t*)|, that *a*_*i*_(*t* + Δ) = (*u*_*i*_, *v*_*i*_, *δ*_*i*_ + Δ).

Under the improper flat prior on the root value, the conditional density of the trait values at *A*^*D*^(*t*), given the observed tip values *y*, is

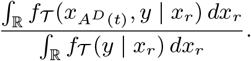

For any tip *s* of *T*, applying Lemma 1 to both the numerator and the denominator shows that this ratio is equal to the conditional density of the trait values at *A*^*D*^(*t*) under the same Brownian model on the rerooted tree *T* ^(*s*)^, given that the trait value at the new root is *y*_*s*_ and given the observed trait values at all the other tips. Let *Y* ^(*s*)^ and *y*^(*s*)^ denote, respectively, the random vector of trait values and the vector of observed trait values, both at the tips of *T* ^(*s*)^, which are the tips of *T* other than *s*. The conditional expectation under the improper flat prior of the empirical mean of the trait values at the positions of *A*^*D*^(*t*) given the tip values of *T*, 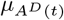, is equal to the conditional expectation, given the observed tip values, of the empirical mean of the trait values at the same positions in the tree *T* ^(*s*)^, namely

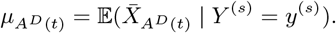

There is at most one index 1 ≤ *k* ≤ |*A*^*D*^(*t*)| such that *s* is a descendant of *v*_*k*_ in *T*. Moreover, there is such an index if and only if there is a position of *A*^*D*^(*t*) on the path from *r* to *s*.

Let us first consider the case where there is no such index *k*. It follows that all the position representations *a*_*i*_(*t*) = (*u*_*i*_, *v*_*i*_, *δ*_*i*_) are such that *u*_*i*_ is closer to *s* than *v*_*i*_ for all 1 ≤ *i* ≤ |*A*^*D*^(*t*)|.

Since the rerooted tree *T*^(*s*)^ is a stem tree with known root value, Lemma 2, applied to the positions of *A*^*D*^(*t*) and *A*^*D*^(*t* + Δ), gives

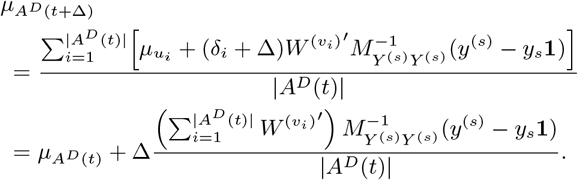

which is affine in Δ.

In the case where there is an index 1 ≤ *k* ≤ |*A*^*D*^(*t*)| such that *s* is a descendant of *v*_*k*_ in *T*, we represent the positions of *A*^*D*^(*t*) in the rerooted tree *T*^(*s*)^ by

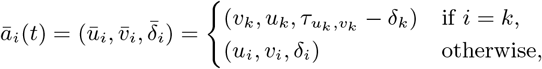

and the positions of *A*^*D*^(*t* + Δ) by

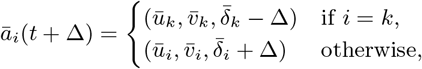

Then, for every 1 ≤ *i* ≤ |*A*^*D*^(*t*)|, *ū*_*i*_ is closer to *s* than 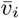. Lemma 2 now gives

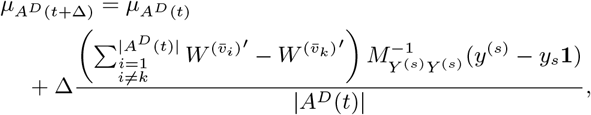

which is affine in Δ.

For the last statement, assume that *T* is ultrametric and that *D* contains all the tips of *T*. For every time *t* under consideration, the sets of tips descending from the nodes *v*_*i*_, for 1 ≤ *i* ≤ |*A*^*D*^(*t*)|, form a partition of the set of tips of *T*. In particular, whatever the tip *s* chosen as the new root, there exists an index *k* such that *s* descends from *v*_*k*_.

Moreover, in the rerooted tree *T*^(*s*)^, the set of tips descending from 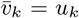 is the union of the sets of tips descending from 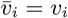, for *i* ≠ *k*. It follows that

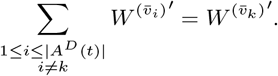

Therefore, the coefficient of Δ in the expression of 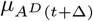 vanishes, and the expected empirical mean is constant between consecutive lineage event times.

## D Proof of Proposition 2

We consider the natural direction of time and assume that the origin of time corresponds to the root *r* of the crown tree *T*. For any time *t*, we write 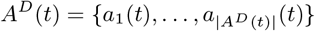. Under the same assumptions and notation as in the preceding section, let *t* be a time in the temporal span of *T* and let Δ ∈ ℝ be such that there is no lineage event time between *t* and *t* + Δ in *T*. It follows that |*A*^*D*^ (*t* + Δ)| = |*A*^*D*^ (*t*)| and *A*^*D*^ (*t*) and *A*^*D*^ (*t* + Δ) can be ordered in a way such that, for all 1 ≤ *i* ≤ |*A*^*D*^ (*t*)|, *a*_*i*_(*t*) = (*u*_*i*_, *v*_*i*_, *δ*_*i*_) and *a*_*i*_(*t* + Δ) = (*u*_*i*_, *v*_*i*_, *δ*_*i*_ + Δ). If |*A*^*D*^(*t*)| = 1, then |*A*^*D*^(*t* + Δ)| = 1 and, by convention, 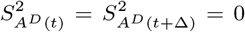. We may therefore assume in what follows that |*A*^*D*^ (*t*)| ≥ 2.

From Lemma 1, the conditional joint distribution of the trait values at *A*^*D*^(*t*) under an improper flat prior for the root of *T*, and in particular its empirical variance over *A*^*D*^(*t*), can equivalently be obtained by rerooting *T* at any of its tips where the trait is known. Let *s* be such a tip and *T*^(*s*)^ be the tree *T* rerooted at *s*. In what follows, all expectations are conditional on the observed tip values, with the trait value at the new root fixed to *y*_*s*_.

We adapt the notations of Section 2.3 and consider

- 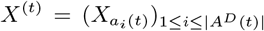 where 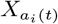 is the random variable associated with the trait value at the position *a*_*i*_ (*t*) of *T*^(*s*)^,
- 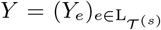 where *Y*_*e*_ is the random variable associated with the trait value at the tip *e* and
- *σ*^2^*M* ^(*t*)^ the variance-covariance matrix of the vector (*X*^(*t*)^, *Y*) with respect to the rerooted tree *T*^(*s*)^.

Under the current assumptions, the covariance matrix *M* ^(*t*+Δ)^ has the same dimension as *M* ^(*t*)^. Although the entries of *M* ^(*t*+Δ)^ are indexed by (*X*^(*t*+Δ)^, *Y*), whereas those of *M* ^(*t*)^ are indexed by (*X*^(*t*)^, *Y*), we identify the index 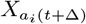 of *M* ^(*t*+Δ)^ with the index 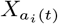 of *M* ^(*t*)^ for every *i*.

For every pair (*U, V*) of coordinates of (*X*^(*t*+Δ)^, *Y*), the entry *M* ^(*t*+Δ)^(*U, V*) is equal to

- *M* ^(*t*)^(*U, V*) + *κ*Δ if there exists 1 ≤ *i* ≤ |*A*^*D*^(*t*)| such that

- either there exists 1 ≤ *j* ≤ |*A*^*D*^(*t*)|, with possibly *i* = *j*, such that *a*_*j*_ (*t*) descends from *a*_*i*_(*t*) in *T* ^(*s*)^ and 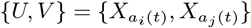,
- or there exists a tip *e* of *T* ^(*s*)^ which descends from *a*_*i*_(*t*) in *T* ^(*s*)^ with 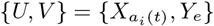,

where

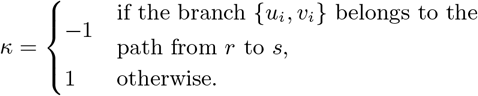

- *M* ^(*t*)^(*U, V*) in all other cases.

Note that 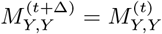. Equation (1) yields

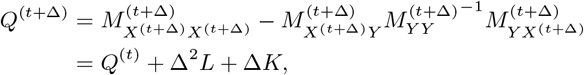

where *L* and *K* are matrices that do not depend on Δ.

Moreover, applying Lemma 2 coordinatewise gives

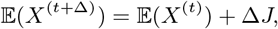

where *J* is a real vector which does not depend on Δ.

From Equation (3), we have

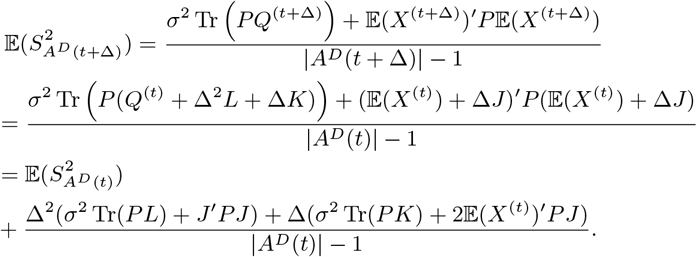

This proves that 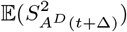 is at most quadratic in Δ.

## Notes

https://github.com/gilles-didier/PastMoments

### Competing Interest Statement

The authors have declared no competing interest.

## References

T.W. Anderson. An Introduction to Multivariate Statistical Analysis. Wiley Series in Probability and Statistics. Wiley-Interscience, Hoboken, NJ, 3rd edition, 2003. ISBN 9780471360919.

Charles N. Ciampaglio, Matthieu Kemp, and Daniel W. McShea. Detecting changes in morphospace occupation patterns in the fossil record: characterization and analysis of measures of disparity. Paleobiology, 27(4):695–715, 2001. doi: 10.1666/0094-8373(2001)027&#60;0695:DCIMOP#x0026;#62;2.0.CO;2.

Noel Cressie and Marinus Borkent. The moment generating function has its moments. Journal of Statistical Planning and Inference, 13:337–344, 1986. ISSN 0378-3758. doi: 10.1016/0378-3758(86)90143-6. URL https://www.sciencedirect.com/science/article/pii/0378375886901436.

Robert B. Davies. The distribution of a linear combination of χ2 random variables. Journal of the Royal Statistical Society Series C: Applied Statistics, 29(3):323–333, 12 1980. ISSN 0035-9254. doi: 10.2307/2346911. URL https://doi.org/10.2307/2346911.

Gilles Didier. Phylogenetic dynamics of MRCA ages and empirical moments of a Brownian trait, 2026. URL https://arxiv.org/abs/2605.29736.

Pierre Duchesne and Pierre Lafaye De Micheaux. Computing the distribution of quadratic forms: Further comparisons between the Liu-Tang-Zhang approximation and exact methods. Computational Statistics and Data Analysis, 54:858–862, 2010.

J. Felsenstein. Phylogenies and the comparative method. Am. Nat., 125:1–15, 1985.

Joseph Felsenstein. Inferring Phylogenies. Sinauer Associates, Sunderland, Massachusetts, 2004.

Mike Foote. The evolution of morphological diversity. Annual Review of Ecology, Evolution, and Systematics, 28(Volume 28, 1997):129–152, 1997. ISSN 1545-2069. doi: 10.1146/annurev.ecolsys.28.1.129. URL https://www.annualreviews.org/content/journals/10.1146/annurev.ecolsys.28.1.129.

Theodore JR. Garland, Peter E. Midford, and Anthony R. Ives. An introduction to phylogenetically based statistical methods, with a new method for confidence intervals on ancestral values. American Zoologist, 39(2):374–388, 04 1999. ISSN 0003-1569. doi: 10.1093/icb/39.2.374. URL https://doi.org/10.1093/icb/39.2.374.

Thomas Guillerme, Natalie Cooper, Stephen L. Brusatte, Katie E. Davis, Andrew L. Jackson, Sylvain Gerber, Anjali Goswami, Kevin Healy, Melanie J. Hopkins, Marc E. H. Jones, Graeme T. Lloyd, Joseph E. O’Reilly, Abi Pate, Mark N. Puttick, Emily J. Rayfield, Erin E. Saupe, Emma Sherratt, Graham J. Slater, Vera Weisbecker, Gavin H. Thomas, and Philip C. J. Donoghue. Disparities in the analysis of morphological disparity. Biology Letters, 16(7): 20200199, 07 2020. ISSN 1744-9561. doi: 10.1098/rsbl.2020.0199. URL https://doi.org/10.1098/rsbl.2020.0199.

Luke J. Harmon, James A. Schulte, Allan Larson, and Jonathan B. Losos. Tempo and mode of evolutionary radiation in iguanian lizards. Science, 301(5635):961–964, 2003. doi: 10.1126/science.1084786. URL https://www.science.org/doi/abs/10.1126/science.1084786.

David A. Harville. Bayesian inference for variance components using only error contrasts. Biometrika, 61(2):383–385, 08 1974. ISSN 0006-3444. doi: 10.1093/biomet/61.2.383. URL https://doi.org/10.1093/biomet/61.2.383.

J. P. Imhof. Computing the distribution of quadratic forms in normal variables. Biometrika, 48(3–4):419–426, 12 1961. ISSN 0006-3444. doi: 10.1093/biomet/48.3-4.419. URL https://doi.org/10.1093/biomet/48.3-4.419.

Dwueng-Chwuan Jhwueng. Two Gaussian bridge processes for mapping continuous trait evolution along phylogenetic trees. Mathematics, 9(16), 2021. ISSN 2227-7390. doi: 10.3390/math9161998. URL https://www.mdpi.com/2227-7390/9/16/1998.

Dieter Korn, Melanie J. Hopkins, and Sonny A. Walton. Extinction space—a method for the quantification and classification of changes in morphospace across extinction boundaries. Evolution, 67(10): 2795–2810, 2013. doi: 10.1111/evo.12162. URL https://onlinelibrary.wiley.com/doi/abs/10.1111/evo.12162.

Bruce Stagg Martin and Marjorie Gail Weber. Stochastic character mapping of continuous traits on phylogenies. Systematic Biology, page syag031, 03 2026. ISSN 1063-5157. doi: 10.1093/sysbio/syag031. URL https://doi.org/10.1093/sysbio/syag031.

Emilia P. Martins and Thomas F. Hansen. Phylogenies and the comparative method: A general approach to incorporating phylogenetic information into the analysis of interspecific data. The American Naturalist, 149(4):646–667, 1997. doi: 10.1086/286013. URL https://www.journals.uchicago.edu/doi/abs/10.1086/286013.

A. M. Mathai and Serge B. Provost. Quadratic forms in random variables : theory and applications. Statistics, text-books and monographs; v.126. Dekker, New York, 1992. ISBN 0824786912.

Kenneth O. McGraw and S. P. Wong. A common language effect size statistic. Psychological Bulletin, 111(2):361–365, 1992. doi: 10.1037/0033-2909.111.2.361.

Manuela Royer-Carenzi and Gilles Didier. A comparison of ancestral state reconstruction methods for quantitative characters. Journal of Theoretical Biology, 404:126–142, 2016. ISSN 0022-5193. doi: 10.1016/j.jtbi.2016.05.029. URL http://www.sciencedirect.com/science/article/pii/S0022519316301205.

D. Schluter, T. Price, A.O. Mooers, and D. Ludwig. Likelihood of ancestor states in adaptive radiation. Evolution, 51(6):1699–1711, 1997.

Graham J. Slater. Phylogenetic evidence for a shift in the mode of mammalian body size evolution at the cretaceous-palaeogene boundary. Methods in Ecology and Evolution, 4(8):734–744, 2013. 10.1111/2041-210X.12084. https://besjournals.onlinelibrary.wiley.com/doi/abs/10.1111/2041-210X.12084. doi: URL

Graham J. Slater, Samantha A. Price, Francesco Santini, and Michael E. Alfaro. Diversity versus disparity and the radiation of modern cetaceans. Proceedings of the Royal Society of London B: Biological Sciences, 2010. ISSN 0962-8452. doi: 10.1098/rspb.2010.0408. URL http://rspb.royalsocietypublishing.org/content/early/2010/05/18/rspb.2010.0408.

András Vargha and Harold D. Delaney. A critique and improvement of the CL common language effect size statistics of McGraw and Wong. Journal of Educational and Behavioral Statistics, 25(2):101–132, 2000. doi: 10.3102/10769986025002101.

Matthew A. Wills, Derek E. G. Briggs, and Richard A. Fortey. Disparity as an evolutionary index: a comparison of cambrian and recent arthropods. Paleobiology, 20 (2):93–130, 1994. doi: 10.1017/S009483730001263X.

